# Preprocessing homologous regions in annotated protein sequences concerning machine-learning applications

**DOI:** 10.1101/2024.10.25.620288

**Authors:** Nawar Malhis

## Abstract

Accurate preprocessing of annotated protein sequences with regard to homologies is essential for maintaining the integrity of machine-learning applications. This study presents two new tools—HAM (Homology-based Annotation Masking) and HAC (Homology Annotation Conflict)— designed to address these challenges. HAM detects and masks homologous regions between datasets to prevent leakage, while HAC identifies and resolves annotation inconsistencies within datasets. Applying these tools to three benchmark datasets revealed substantial overlooked homology and annotation conflicts, even in datasets that had been previously clustered by sequence identity. These findings underscore the importance of homology-aware preprocessing to ensure the integrity of model training and evaluation. By integrating HAM and HAC into machine learning workflows, researchers can improve the consistency and trustworthiness of protein sequence-based predictions.

**Availability:** github.com/NawarMalhis/HAM.git

## 1 Introduction

Machine learning tools are extensively used to predict various functional features of protein sequences, including antimicrobial peptides (AMPs) [1], antigen-antibody affinity [2], post-translational modification (PTM) sites [3], and protein-disordered binding regions [4][5]. These models are trained using protein sequences annotated with the target feature, which are typically divided into separate training and testing datasets. The training set is used to build the model, while the testing set is used to evaluate its predictive performance.

Two critical issues must be addressed to ensure the integrity of this process. First is **data leakage** between the training and testing sets. In theory, training and testing datasets should be separated at random, however, this strategy can introduce data leakage in the context of protein sequences due to the high prevalence of homology. A common solution is to cluster sequences based on sequence identity and then randomly split the clusters, ensuring that homologous sequences reside exclusively in either the training or testing set. The identity cut-off used for clustering is typically tailored to the functional feature under investigation. For features localized to short sequence regions, a stricter (lower) identity threshold is required to minimize homology-driven leakage relevant to the feature.

The second issue is **annotation conflict**. Regions with a high degree of homology tend to share similar functions and, crucially, are likely to receive similar predictions. If conflicting annotations exist among such highly homologous regions in the training set, the model may learn inconsistent signals. Similarly, conflicts in the test set can impair the reliability of model evaluation.

In this study, I introduce two tools designed to detect and mitigate these issues in annotated protein sequence datasets. The first, **HAM (<u>H</u>omology-based <u>A</u>nnotation <u>M</u>asking)**, addresses data leakage by identifying homologous regions across datasets, detecting cross-annotations, and masking overlapping regions in one of the datasets. The second tool, **HAC (<u>H</u>omology <u>A</u>nnotation <u>C</u>onflict)**, operates within a single dataset to identify local homologies with conflicting annotations and resolves these conflicts to improve consistency.

## 2 Results

I examined three benchmark datasets, each split into training and testing sets, designed for developing and evaluating predictors of intrinsically disordered protein binding sites. The first dataset, **D2008**, was compiled in 2008 [6] and has since been widely used to train and evaluate multiple MoRF (Molecular Recognition Feature) predictors, including MoRFpred [6], MFSPSSMpred [7], MoRFchibi [8], MoRFchibi_web [9], fMoRFpred [10], MoRFchibi_light [11], OPAL [12], MoRFmlp [13], and MoRFcnn [14]. The second dataset, **D2021**, was assembled in 2021 and used to develop and assess DeepDISObind, a predictor of binding sites within intrinsically disordered regions (IDRs) [15]. The third dataset, **D2022**, was introduced in 2022 for training and testing CLIP, a method designed to predict linear interacting peptides (LIPs) [16].

The authors of MoRFpred assembled a dataset of 840 protein sequences containing intrinsically disordered regions (IDRs) that bind to protein targets, known as molecular recognition features (MoRFs), which undergo disorder-to-order transitions upon binding. This dataset, referred to as D2008, was clustered at 30% sequence identity using CD-HIT [17], resulting in 427 clusters. These clusters were then randomly divided into a training set of 421 sequences and a testing set of 419 sequences.

The authors of DeepDISOBind compiled a separate dataset, D2021, consisting of 750 protein sequences from the DisProt database [18], each containing high-quality annotations of IDR binding sites that interact with proteins, DNA, or RNA. For this analysis, only protein-binding IDR sites are considered. Like D2008, the sequences were clustered at 30% identity using CD-HIT, producing 650 clusters. These were randomly split into a training set of 356 sequences and a testing set of 394 sequences. Notably, despite having fewer total sequences than D2008, D2021 produced more clusters, indicating a lower degree of sequence homology within the dataset.

The creators of CLIP began with over 1,800 sequences annotated with Linear Interacting Peptides (LIPs) from the MobiDB 3.0 LIP database [19]. Initially, they clustered sequences at 100% identity using CD-HIT, retaining only the longest sequence from each cluster while transferring LIP annotations from other members within the same cluster. These sequences were then reclustered at a 25% identity threshold, yielding 1,440 clusters. Again, the longest sequence from each cluster was selected, and the rest were discarded, forming the D2022 dataset. Finally, the resulting sequences were randomly divided into a training set of 1,000 sequences and a testing set of 440 sequences.

In summary, the limited number of known MoRFs at the time D2008 was constructed led the MoRFpred authors to retain homologous sequences to ensure dataset size. In contrast, D2021 contains fewer sequences but greater sequence diversity, resulting in lower homology. The extensive annotation coverage in the MobiDB LIP database enabled the authors of CLIP to construct a much larger dataset (D2022) and apply stringent homology filtering at a 25% identity threshold, both within and between the training and testing sets.

### 2.1 Identifying and addressing data leakage between training and testing sets using HAM

Despite applying clustering strategies to partition each of the three datasets into training and testing sets, the use of HAM revealed significant instances of data leakage. HAM identified notable homology between annotated regions in the training and testing sequences across all three datasets. **Table 1** summarizes the extent of this homology-based overlap for each dataset. It is important to note that the overlap values from Dataset A to Dataset B may differ from those in the reverse direction, as a single residue in one dataset may be homologous to multiple residues in the other (see **Figure 1** for illustration).

**Table 1:** The table reports the homology overlap between annotated sequences in the training and testing datasets, as identified by HAM using its default parameters. The identity cut-off indicates the sequence identity threshold used during clustering to separate the datasets. Total refers to the number of homologous residues found in Dataset 1. The entry “A1 to A2” represents the number of residues annotated with feature A1 in Dataset 1 that are homologous to residues annotated with feature A2 in Dataset 2.

| Dataset 1 | Dataset 2 | Identity cut-off | Total | 0 to 0 | 0 to 1 | 1 to 0 | 1 to 1 |
| --- | --- | --- | --- | --- | --- | --- | --- |
| TR2008 | TS2008 | 30% | 46 | 37 | 2 | 1 | 6 |
| TS2008 | TR2008 | 30% | 76 | 61 | 1 | 3 | 11 |
| TR2021 | TS2021 | 30% | 761 | 739 | 11 | 11 | 0 |
| TS2021 | TR2021 | 30% | 702 | 680 | 11 | 11 | 0 |
| TR2022 | TS2022 | 25% | 375 | 368 | 1 | 6 | 0 |
| TS2022 | TR2022 | 25% | 375 | 368 | 6 | 1 | 0 |

**Figure 1:**
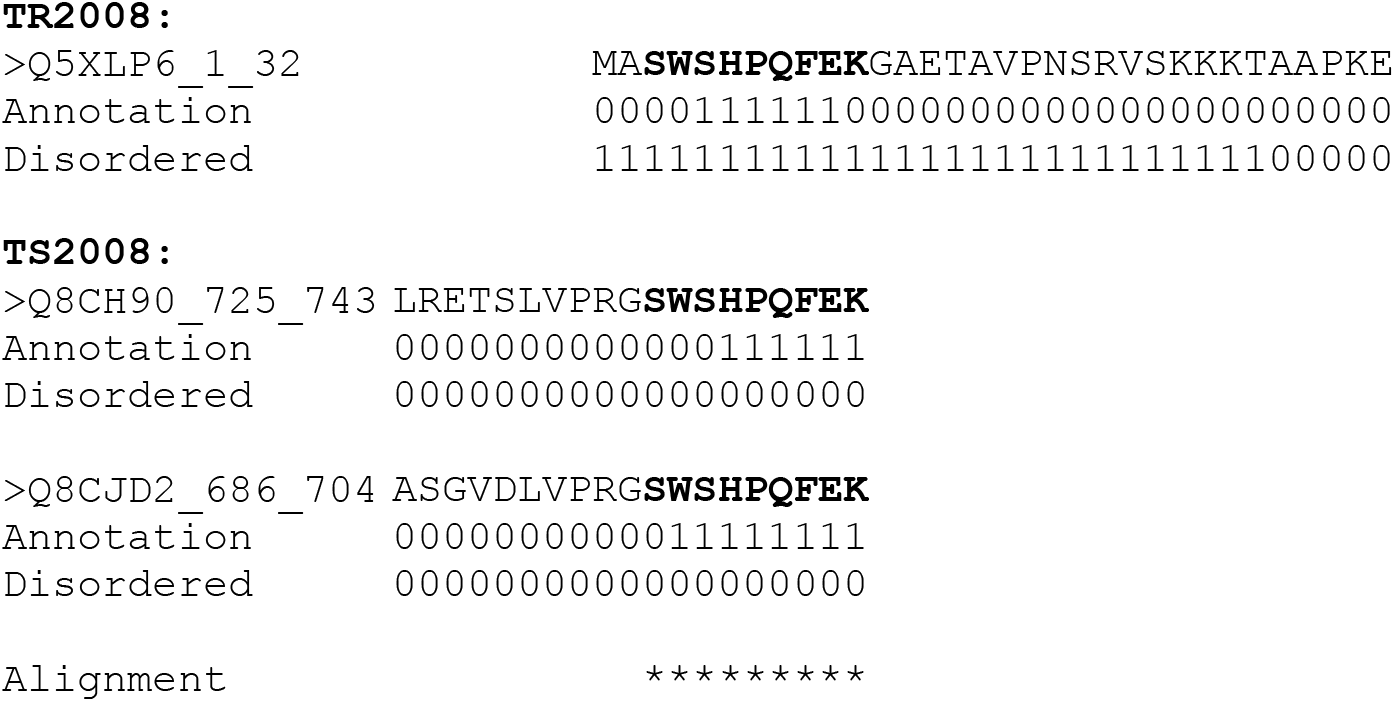
Although the training (TR2008) and testing (TS2008) datasets were separated using a 30% sequence identity cut-off, HAM identified a nine-residue region with 100% identity shared between the N-terminus of the training sequence Q5XLP6 (1kl3_E) and the C-terminus of two testing sequences, Q8CH90 (1kl3_F) and Q8CJD2 (1rsu_P). In the training sequence Q5XLP6, six of these nine residues are annotated as MoRFs. In the testing sequences, six residues are annotated as MoRFs in Q8CH90 and eight in Q8CJD2.

The following example highlights a case of conflicting annotations for homologous residues between the D2021 training and testing datasets. Although the overall sequence identity between P63315 and P16471 is only 5.3%, HAM identified an eleven-residue region with high local identity (9 out of 11 identical residues) shared between P63315 (residues 90–100) from the TR2021 training set and P16471 (residues 286–296) from the TS2021 testing set. In P16471, these residues are annotated as IDR-binding, while in P63315, they are annotated as non-binding. Interestingly, MoRFchibi_web predictions indicate that residues 90–100 in P63315 are likely to be IDR-binding (see **Figure 2**).

**Figure 2:**
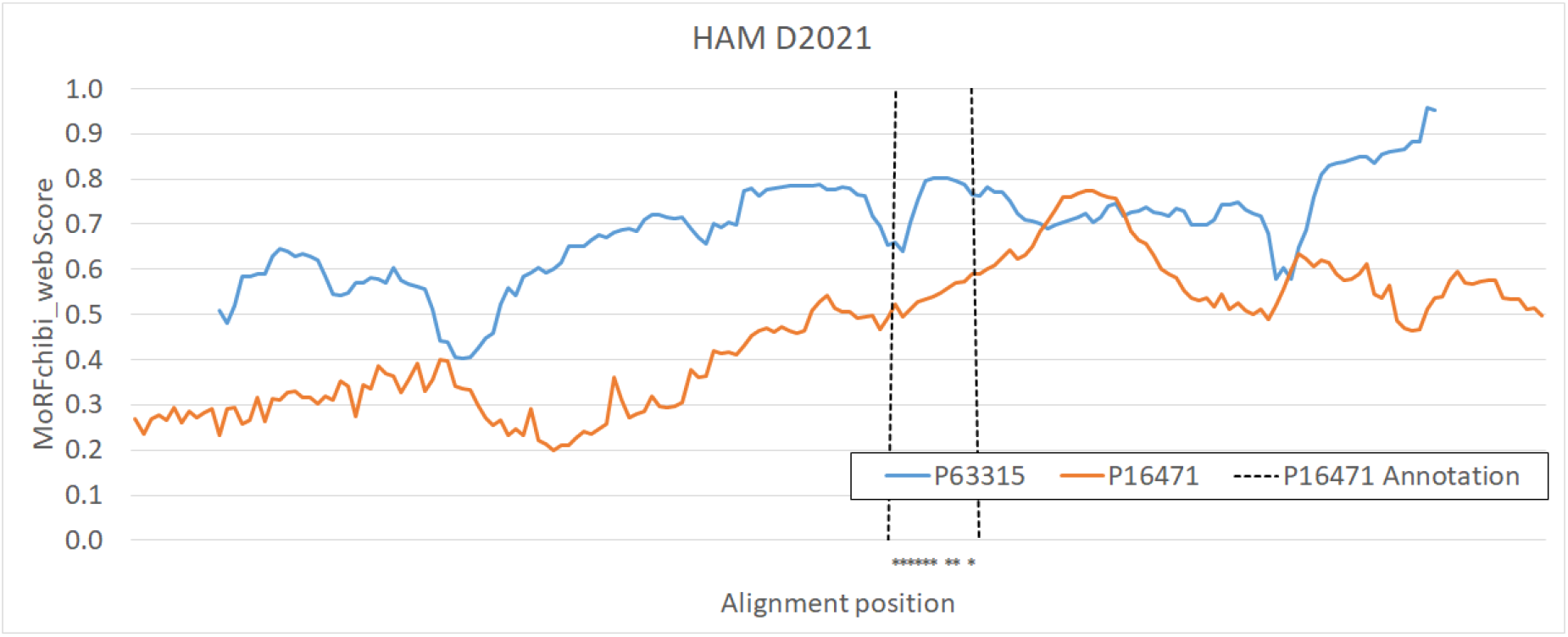
MoRFchibi_web scores for P63315 and the corresponding region of P16471 (residues 186–371) are shown. The aligned region between the two sequences is indicated by asterisks (*). Although this region is not annotated as a protein-binding site in the training sequence (P63315), it is annotated as such in the testing sequence (P16471). Notably, MoRFchibi_web predicts the region in P63315 as a protein-binding MoRF.

In the D2022 dataset, HAM identified the C-terminal 98 residues of the training protein Q13351 (KLF1_HUMAN, 362 residues) as homologous to the corresponding region in the testing protein Q60793 (KLF4_MOUSE, 483 residues), with a sequence identity of 83.7%. Although both regions are annotated as non-binding, this high similarity constitutes data leakage between the training dataset (TR2022) and the testing dataset (TS2022). HAM detects such homologous regions and provides the option to mask them in one of the datasets to prevent contamination.

### 2.1 Using HAC for identifying and processing annotation conflicts within sequences in the same dataset

Unlike HAM, which focuses on homology between separate datasets, the Homology Annotation Conflict (HAC) software identifies homologous regions within the same dataset, including those with conflicting annotations. **Table 2** summarizes the annotated homology detected by HAC across the six datasets, using its default parameters.

**Table 2:** Using the HAC software with default parameters, the annotated homology within each of the six datasets. The Identity cut-off is the clustering cut-off used, if any. Total is the total homologous residues. “A1 to A2” is the number of residues annotated with A1 that are homologous to residues annotated with A2.

| Dataset | Identity cut-off | Total | 0 to 0 | 0 to 1 | 1 to 0 | 1 to 1 |
| --- | --- | --- | --- | --- | --- | --- |
| TR2008 | None | 89,309 | 84,278 | 1,652 | 1,092 | 2,287 |
| TS2008 | None | 113,986 | 109,404 | 2,064 | 1,102 | 1,416 |
| TR2021 | None | 3,938 | 2,840 | 353 | 353 | 392 |
| TS2021 | None | 4,464 | 3,849 | 302 | 313 | 0 |
| TR2022 | 25% | 1,125 | 1,067 | 13 | 13 | 32 |
| TS2022 | 25% | 310 | 310 | 0 | 0 | 0 |

Figure 3. illustrates an annotation conflict within a 13-residue homologous region shared among three large ribosomal subunit protein sequences in the TS2021 dataset: P05318 (residues 94–106), P05387 (residues 103–115), and P23632 (residues 95–107). This shared region lies within a broader intrinsically disordered region (IDR) annotated in the UniProt database [20] and identified in all three sequences through SAM-based analysis using MobiDB-lite [19].

Figure 4. displays seven homologous sequences with annotation conflicts identified by HAC in the TR2008 dataset, along with three resolution strategies provided by the `hac_resolve_conflict.py`:

- Conflicting to 1 – all conflicting annotations are resolved as ‘1’;
- Conflicting to 0 – all conflicts are resolved as ‘0’;
- Conflicting to mask – conflicting annotations are masked, i.e., replaced with ‘-’ (unknown).

**Figure 3:**
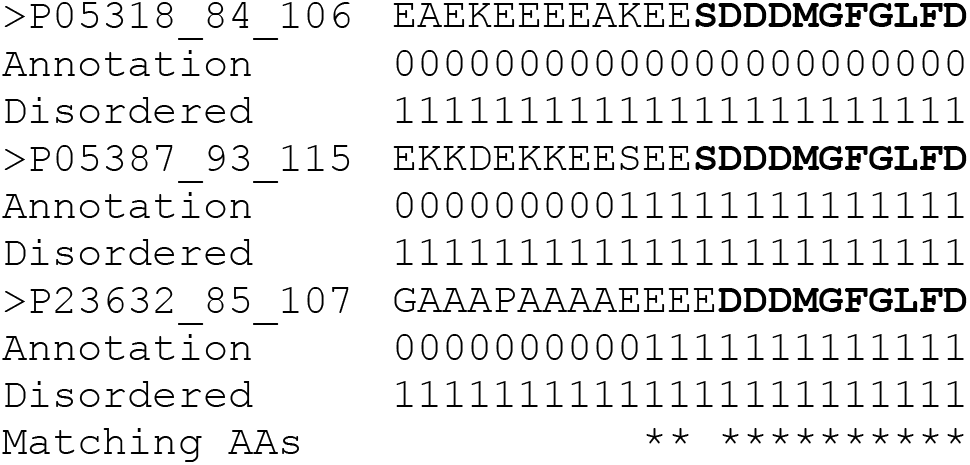
The aligned disordered C-terminal regions of three large ribosomal subunit proteins from the TS2021 dataset—P05318 (Yeast), P05387 (Human), and P23632 (Trypanosoma cruzi)—share a 13-residue homologous segment with at least 12 out of 13 residues identical between each pair. This region is annotated as protein-binding in P05387 and P23632, but as non-binding in P05318, highlighting an annotation conflict.

**Figure 4:**
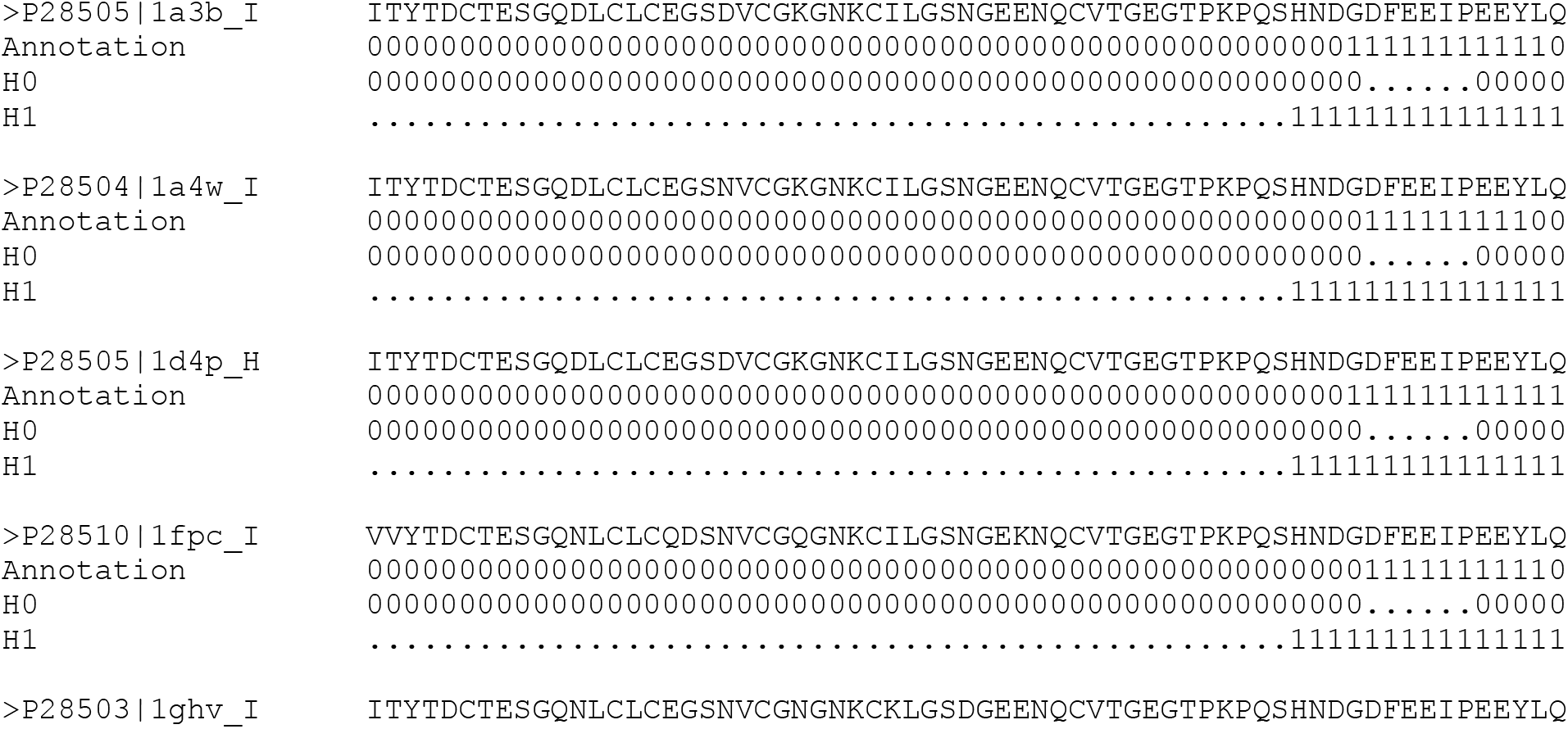
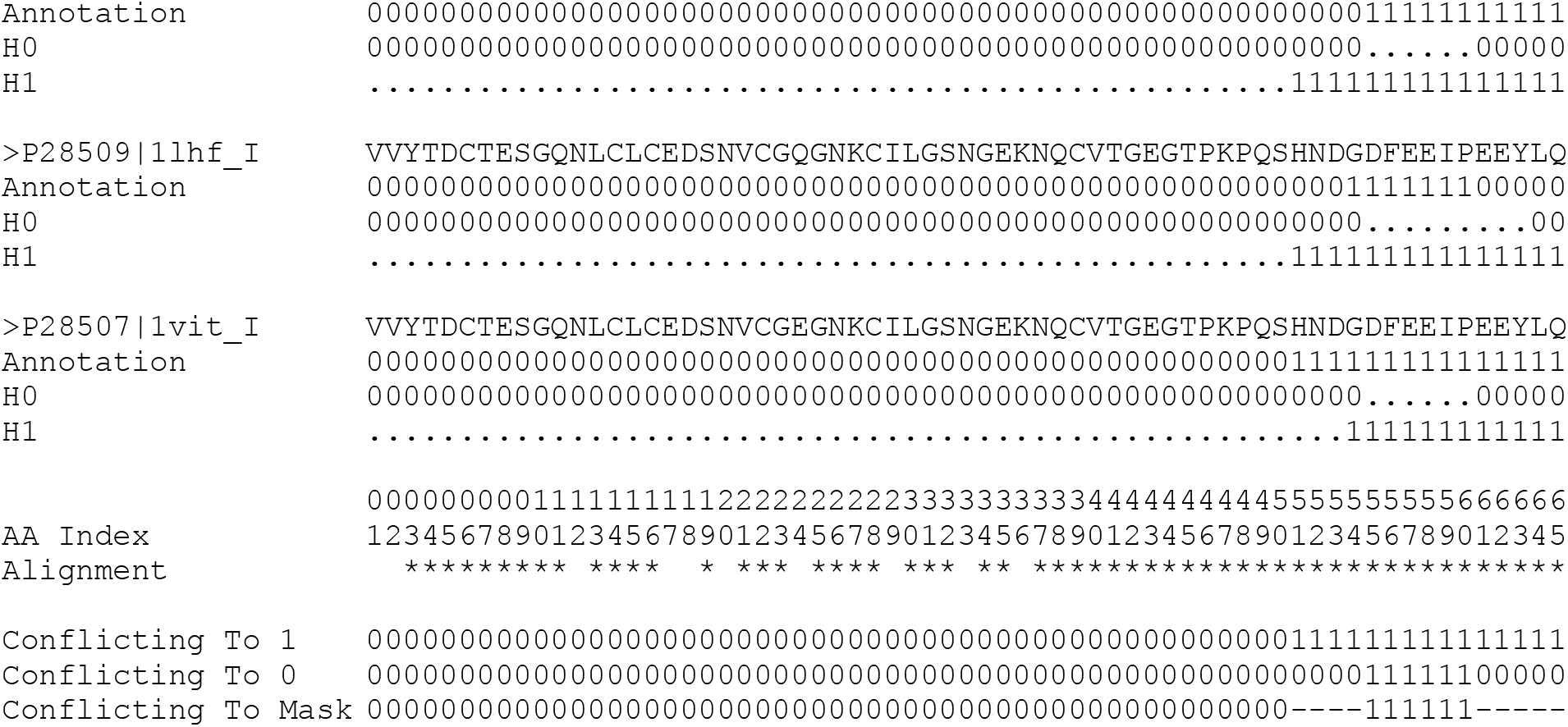
Seven aligned, equal-length annotated sequences from the TR2008 dataset with conflicting annotations are shown. H0 indicates positions annotated as '0' in homologous sequences, and H1 indicates positions annotated as '1'. The final three lines illustrate the conflict resolution strategies provided by `hac_resolve_conflict.py`: '0' to '1', '1' to '0', and the masked out option.

HAC also detected extended regions of homology between annotated sequences. In the TS2021 dataset, HAC identified two highly similar proteins: P05204 (a 90-residue non-histone chromosomal protein, HMG-17, from Homo sapiens) and P02313 (its 90-residue homolog from Bos taurus), which share 97.8% sequence identity. Despite this similarity, there is a clear annotation conflict—P05204 is entirely annotated as protein-binding, whereas P02313 is annotated as non-binding. According to UniProt, both proteins are predicted to be disordered based on automatic assertions from MobiDB-lite.

Another example from the TR2021 dataset involves residues 1–77 of P0ABD8 and P0ABE0, both biotin carboxyl carrier proteins of acetyl-CoA carboxylase from E. coli strains K12 and O157:H7, respectively. These proteins are identical in length (156 residues) and share 100% sequence identity. However, residues 1–77 are annotated as protein-binding in P0ABD8, while P0ABE0 is annotated as entirely non-binding, highlighting a significant annotation conflict.

## 3. Methods

The software package consists of four Python tools: `ham.py`, `ham_mask_homology.py`, `hac.py`, and `hac_resolve_conflicts.py`.

Both `ham.py` and `hac.py` use `BLASTP` with the parameters `-outfmt 6` and `-word_size 3` to identify all possible local alignments between a query sequence and a target database.

- In `ham.py`, the database is built from the larger of the two input datasets, while the sequences from the other dataset serve as queries.
- In `hac.py`, the same input dataset is used for both the database and the queries, with self-matches excluded.

In both tools, matches are retained if they contain no gaps, are longer than a minimum threshold (default: 10 amino acids), and have a sequence identity above a specified cut-off (default: 80%).

### 3.1 Output Files

`ham.py` produces three outputs:

- A detailed file TSV (Tab-Separated Values) with one-to-one residue alignment matches.
- Two annotated FASTA files (one for each input dataset), including additional annotation lines that reflect homologous residue annotations.

`ham_mask_homology.py` reads all annotated FASTA files generated by `ham.py` for a given dataset and outputs a new annotated FASTA file with all homologous residues masked—i.e., annotated with ‘-’.

`hac.py` generates:

- A detailed alignment file TSV with one-to-one homologous residue matches.
- An annotated FASTA file that includes homologous annotation information for each sequence.

`hac_resolve_conflicts.py` processes the annotated FASTA file produced by hac.py, resolving any annotation conflicts by reannotating each conflicting residue as ‘1’, ‘0’, or masking it with ‘-’, according to the selected resolution strategy (see **Figure 4**).

## 4. Discussion

Accurate preprocessing of protein sequence datasets is a critical prerequisite for the successful application of machine learning methods to predict functional features such as intrinsically disordered protein binding sites. This study highlights two major challenges often overlooked in current workflows: data leakage resulting from homologous regions shared between training and testing datasets, and annotation conflicts within homologous regions in the same dataset. Both issues have important implications for model training fidelity and evaluation accuracy.

Our results demonstrate that even when standard clustering methods are applied at common identity thresholds (e.g., 30% or 25%) to separate training and testing sets, significant homology-based overlap persists. The use of HAM revealed such overlaps across all three benchmark datasets examined (D2008, D2021, and D2022), uncovering homologous residues shared between datasets that can lead to inadvertent data leakage. This finding underscores that clustering alone, particularly when based on global sequence identity cut-offs, may not be sufficient to prevent local homologous regions from contaminating the independence of training and testing data. Furthermore, the identification of homologous regions with conflicting annotations between datasets (e.g., binding vs. non-binding) adds another layer of complexity that may confound model evaluation and interpretation.

Within datasets, HAC exposed pervasive annotation conflicts in homologous regions that can undermine the internal consistency necessary for effective machine learning. Our examples involving proteins with near-identical sequences yet divergent annotations highlight the potential for misleading signals during model training, which can degrade predictive performance or yield contradictory outcomes. Configurable conflict resolution strategies within HAC allow users to tailor preprocessing to their specific use case.

The application of HAM and HAC to diverse and widely used datasets shows their broad utility and the benefit of integrating these tools into routine preprocessing pipelines. Particularly for features localized to short sequence segments—such as molecular recognition features or linear interacting peptides— careful handling of homology and annotation consistency is vital to avoid overestimating model generalization.

In conclusion, HAM and HAC address critical gaps in preprocessing protein sequence data by systematically detecting and mitigating homology-induced data leakage and annotation conflicts. Incorporating these tools into machine learning workflows will enhance the reliability and interpretability of functional predictions derived from protein sequences, ultimately advancing computational biology applications.

